# Subthalamic nucleus and sensorimotor cortex activity during speech production

**DOI:** 10.1101/463141

**Authors:** A Chrabaszcz, WJ Neumann, O Stretcu, WJ Lipski, A Bush, C Dastolfo-Hromack, D Wang, DJ Crammond, S Shaiman, MW Dickey, LL Holt, RS Turner, JA Fiez, RM Richardson

**Affiliations:** Department of Psychology, University of Pittsburgh, 15213; Movement Disorder and Neuromodulation Unit, Department of Neurology, Campus Mitte, Charité – Universitätsmedizin Berlin, Berlin, Germany, 10117; Machine Learning Department, School of Computer Science, Carnegie Mellon University, 15213; Brain Modulation Lab, Department of Neurological Surgery, University of Pittsburgh School of Medicine, 15213; Department of Physics, FCEN, University of Buenos Aires and IFIBA-CONICET, Argentina, 1428; School of Medicine, Tsinghua University, Beijing, China 100084; Department of Communication Science and Disorders, University of Pittsburgh, 15213; Department of Psychology, Carnegie Mellon University, 15213; Department of Neurobiology, University of Pittsburgh School of Medicine, 15213; University of Pittsburgh Brain Institute, 15213

**Keywords:** speech, vocal tract articulators, subthalamic nucleus, sensorimotor cortex, high gamma oscillations, electrocorticography, deep brain stimulation, Parkinson’s disease

## Abstract

The sensorimotor cortex is somatotopically organized to represent the vocal tract articulators, such as lips, tongue, larynx, and jaw. How speech and articulatory features are encoded at the subcortical level, however, remains largely unknown. We analyzed local field potential (LFP) recordings from the subthalamic nucleus (STN) and simultaneous electrocorticography recordings from the sensorimotor cortex of 11 patients (1 female) with Parkinson’s disease during implantation of deep brain stimulation (DBS) electrodes, while patients read aloud three-phoneme words. The initial phonemes involved either articulation primarily with the tongue (coronal consonants) or the lips (labial consonants). We observed significant increases in high gamma (60–150 Hz) power in both the STN and the sensorimotor cortex that began before speech onset and persisted for the duration of speech articulation. As expected from previous reports, in the sensorimotor cortex, the primary articulator involved in the production of the initial consonant was topographically represented by high gamma activity. We found that STN high gamma activity also demonstrated specificity for the primary articulator, although no clear topography was observed. In general, subthalamic high gamma activity varied along the ventral-dorsal trajectory of the electrodes, with greater high gamma power recorded in the more dorsal locations of the STN. These results demonstrate that articulator-specific speech information is contained within high gamma activity of the STN, with similar temporal but less specific topographical organization, compared to similar information encoded in the sensorimotor cortex.

**SIGNIFICANCE STATEMENT:** Clinical and electrophysiological evidence suggests that the subthalamic nucleus is involved in speech, however, this important basal ganglia node is ignored in current models of speech production. We previously showed that subthalamic nucleus neurons differentially encode early and late aspects of speech production, but no previous studies have examined subthalamic functional organization for speech articulators. Using simultaneous local field potential recordings from the sensorimotor cortex and the subthalamic nucleus in patients with Parkinson’s disease undergoing deep brain stimulation surgery, we discovered that subthalamic nucleus high gamma activity tracks speech production at the level of vocal tract articulators, with high gamma power beginning to increase prior to the onset of vocalization, similar to cortical articulatory encoding.

## INTRODUCTION

Speech articulation constitutes a complex motor behavior involving a precise coordination of different parts of the vocal apparatus, known as articulators (e.g., lips, tongue). While recruitment of the cortical regions in the articulatory realization of speech is widely documented, the specific contributions of different subcortical structures remain largely unknown. Here, for the first time, we use local field potential (LFP) recordings from the subthalamic nucleus (STN) and simultaneous electrocorticography (ECoG) recordings from the sensorimotor cortex to investigate the role of the STN in speech articulation and to compare its spatial and temporal organization for encoding of speech articulators with that of the sensorimotor cortex.

Ample evidence has implicated the ventral-lateral orofacial area of the sensorimotor cortex as a principal cortical region for the neural representation of speech articulators. Electrical stimulation of this region produces somatotopically organized sensorimotor responses for the larynx, tongue, jaw, and lips along the ventral-to-dorsal orientation of the central sulcus, respectively (Penfield and Boldrey, 1937; Penfield, 1954; Woolsey, Erickson, and Gilson, 1979; Breshears, Molinaro, and Chang, 2015). Functional imaging (fMRI) studies generally provide corroborating evidence for the somatotopic cortical representation of the vocal tract effectors, among other body parts, albeit with a varying degree of overlap among individuals (Lotze et al., 2000; Hesselmann et al., 2004; Pulvermüller et al., 2006; Brown, Ngan, and Liotti, 2007; Meier et al., 2008; Takai, Brown, and Liotti, 2010; Carey et al., 2017). Recently, ECoG studies have elaborated the notion of cortical articulatory somatotopy by revealing differentiated neural representations for fine-grained phonetic features and complex kinematics underlying speech articulation (Bouchard et al., 2013; Bouchard & Chang, 2014; Mugler et al., 2014; Lotte et al., 2015; Bouchard et al., 2016; Cheung et al., 2016; Ramsey et al., 2017; Chartier et al., 2018; Conant et al., 2018).

Anatomical connections between the sensorimotor cortex and the basal ganglia via a cortico-striatal-thalamic loop (Alexander, DeLong, and Strick, 1986) suggest that the basal ganglia, including the STN, may also participate in speech production. Indeed, indirect evidence from lesion literature (Brunner et al., 1982; Damasio et al., 1982; Wallesch et al., 1983; Nadeau and Crosson, 1997), from clinical data on deep brain stimulation (DBS) outcomes (Morrison et al., 2004; Witt et al., 2008; Aldridge et al., 2016; Knowles et al., 2018) and neurological disorders involving the basal ganglia (Logemann et al., 1978; Ho et al., 1998; Walsh and Smith, 2012) implicates the basal ganglia in many aspects of speech production. Direct evidence from electrophysiological recordings of STN activity during speech production shows desynchronization of beta power during articulation of non-propositional speech (Hebb, Darvas, and Miller, 2012), and speech-related changes in single unit firing activity (Watson and Montgomery, 2006; Lipski et al., 2018). To our knowledge, however, no study has investigated the spatial and temporal distribution of speech-related neuronal activity for different articulators in the STN relative to the sensorimotor cortex. Given that the STN is anatomically subdivided into sensorimotor, limbic, and associative functional areas (Hamani et al., 2004; Temel et al., 2005; Haynes and Haber, 2013) and that a somatotopic organization for arms, legs, eyes and face is observed within the motor territory of the STN in human and non-human primates (Monakow, Akert, and Kiinzle, 1978; DeLong, Crutcher, and Georgopoulos, 1985; Wichmann, Bergman, and DeLong, 1994; Rodriguez-Oroz et al., 2001; Starr, Theodosopoulos, and Turner, 2003; Theodosopoulos et al., 2003; Nambu, 2011), it is possible that a functional somatotopy for the vocal tract articulators is also maintained within the STN.

We employed a novel experimental paradigm in awake, speaking patients undergoing STN-DBS for Parkinson’s disease, where sensorimotor electrocorticography is recorded simultaneously with STN LFPs. We discovered that STN high gamma (60–150 Hz) activity is dynamic during the production of speech, exhibiting activity that tracks with specific articulatory motor features. Our data further suggest that spatial and temporal characteristics of the neural representations of speech articulators may differ between the cortex and STN.

## MATERIALS AND METHODS

### Participants

Participants included 11 native English-speaking patients with Parkinson’s disease (10M/1F, age: 67.5±7.7 years, duration of disease: 8±2.4 years) undergoing awake stereotactic neurosurgery for implantation of DBS electrodes in the STN. In addition to the clinical subcortical mapping and as part of an IRB approved research protocol, participants were temporarily implanted with subdural electrode arrays over the left ventral sensorimotor cortex. All patients completed Unified Parkinson’s Disease Rating Scale (UPDRS) testing within four months before the surgery. Dopaminergic medication was withdrawn the night before surgery. Subjects’ demographic and clinical characteristics are provided in Table 1. All procedures were approved by the University of Pittsburgh Institutional Review Board (IRB Protocol # PRO13110420), and all patients provided informed consent to participate in the study.

**Table 1.** Subject demographic and clinical characteristics.

| Subject | Gender | Age | Handedness | Education, years | Duration of disease, years | Hoehn and Yahr Stage | UPDRS Score (off medication) |
| --- | --- | --- | --- | --- | --- | --- | --- |
| 1 | Male | 71 | not recorded | not recorded | 6 | 2 | 35 |
| 2 | Male | 60 | Right | 12 | 14 | 2 | 53 |
| 3 | Male | 69 | Right | 14 | 9 | 2 | 46 |
| 4 | Male | 61 | Right | 16 | 5 | 2 | 31 |
| 5 | Male | 68 | Left | 16 | 8 | 2 | 50 |
| 6 | Male | 57 | not recorded | not recorded | 7 | 2 | 44 |
| 7 | Male | 82 | Right | 16 | 8 | 2 | 36 |
| 8 | Male | 66 | Right | 19 | 7 | 2 | 45 |
| 9 | Female | 71 | Right | 16 | 8 | 2 | 24 |
| 10 | Male | 77 | Right | 18 | 10 | 2 | 27 |
| 11 | Male | 60 | Right | 13 | 6 | 2 | 39 |

### Stimuli and procedure

Participants performed a reading-aloud task during the subcortical mapping portion of the surgery in up to 4 recording sessions per patient, with 120 trials per session. The visual stimuli consisted of consonant-vowel-consonant (CVC) words and pseudowords presented on a computer screen. The stimuli were chosen from an existing stimulus set, and were balanced along a number of psycholinguistic parameters, such as phonological and orthographic neighborhood density, bigram frequency, phonotactic and biphone probability, etc. (for a detailed description of the stimuli, see Moore, Fiez, and Tompkins, 2017). For the purposes of the present study, the stimuli were grouped into two categories based on the primary articulator involved in the production of the initial consonants: words with word-initial labial consonants (i.e., those requiring closure or constriction of the air flow primarily with the use of the lips), and words with word-initial coronal consonants (i.e., those requiring articulation primarily with the use of the tongue). The labial consonants subsumed bilabial (/p/, /m/) and labiodental (/f/, /v/) phonemes; coronal consonants included alveolar (/s/, /z/, /t/, /d/, /l/, /n/), post-alveolar (/∫/, /r/), and dental (/θ/, /ð/) phonemes.

The stimuli were created and presented by custom code running in the Matlab environment (MathWorks, Natick, MA) using Psychophysics Toolbox extensions (Brainard, 1997). A schematic of the experimental procedure is shown in Figure 1. On each trial, participants were presented with a white cross against a black background during an intertrial interval, after which a green fixation cross appeared on the screen for 250 ms instructing the participants to get ready. It was followed by a variable interstimulus interval (500-1000 ms) during which the screen remained black. Then the stimulus word was presented on the screen and participants were instructed to read it out loud. The stimulus word remained on the screen until participants made the response, after which the experimenter advanced the presentation to the next trial. All stimuli (120 trials per recording session) were pseudorandomized in order of presentation. Participants were familiarized with the task prior to surgery.

**Figure 1.**
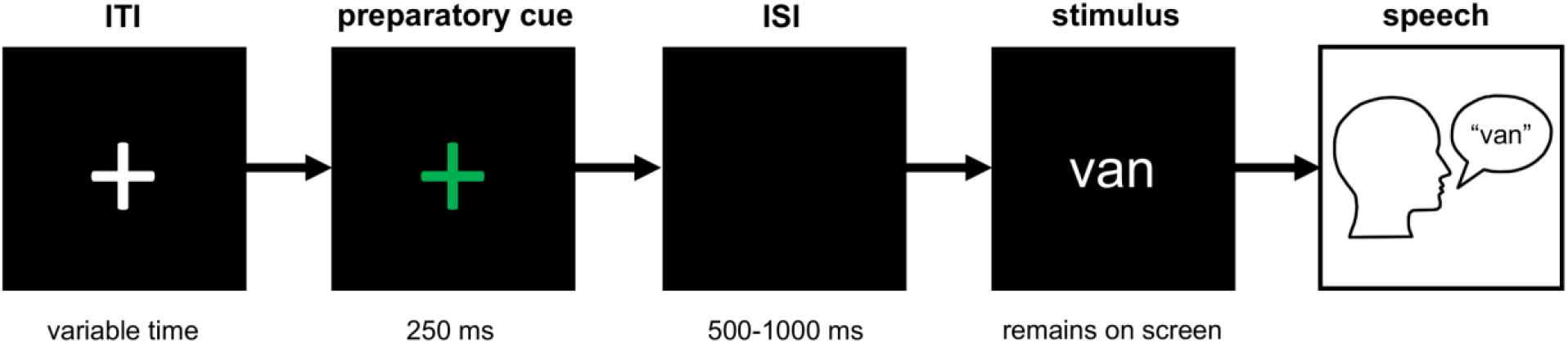
Experimental paradigm. ITI = intertrial interval; ISI = interstimulus interval

### Audio recordings

Participants’ reading aloud was recorded using an omnidirectional microphone (Audio-Technica ATR3350iS Mic, frequency response 50-18,000 Hz, or PreSonus PRM1 Mic, frequency response 20-20,000 Hz). The microphone was positioned at a distance of approximately 8 cm from the subject’s left oral angle of the mouth and oriented at an angle of approximately 45 degrees. A Zoom H6 digital recorder was used to record the audio signal at a sampling rate of 96 kHz. This signal was simultaneously recorded using a Grapevine Neural Interface Processor (Ripple LLC, Salt Lake City, UT, USA) at a lower sampling rate of 30 kHz. The audio recordings were segmented and transcribed offline by phonetically-trained communication science students using the International Phonetic Alphabet (IPA) in a custom-designed graphical user interface (GUI) implemented in MATLAB. The audio recordings were synchronized with the neural recordings using digital pulses delivered to the Neuro-Omega system (Alpha Omega, Nazareth, Israel) via a USB data acquisition unit (Measurement Computing, Norton, MA, model USB-1208FS).

### Subthalamic nucleus recordings

Subjects were implanted with DBS leads bilaterally, but local field potentials were recorded during the administration of the reading-aloud task only for the left side surgery (see Figure 2A for an example of lead trajectory). The LFP signal was acquired with the Neuro-Omega recording system using parylene insulated tungsten microelectrodes (25 μm in diameter, 100 μm in length) with a stainless steel macroelectrode ring (0.55 mm in diameter, 1.4 mm in length) 3 mm above the tip of the microelectrode. The LFP signal was recorded at a sampling rate of 44 kHz and was band-pass filtered at 0.075 Hz to 10 kHz. The microelectrodes targeted the dorsolateral area of the STN, as previously described in Lee et al. (2018). The microelectrodes were oriented on the microtargeting drive system using two or three trajectories of a standard cross-shaped Ben-Gun array with a 2 mm center-to-center spacing: for mapping of the center, posterior, and medial tracts. The microelectrodes were advanced manually in 0.1 mm steps starting 10 mm above the defined target. The patients were subsequently implanted with DBS Medtronic 3389 leads with four platinum-iridium cylindrical macroelectrodes 1.27 mm in diameter, 1.5 mm in length and a 0.5 mm electrode spacing (Medtronic, Minneapolis, MN, USA). The superior and inferior boundaries of the STN were determined by the neurophysiologist and neurosurgeon based on the characteristic STN single-unit neuronal activity obtained from the microelectrode recordings (MER). The speech task was administered and LFP data acquired for up to four different depths within the STN per patient. As a result, LFP data from a total of 79 recording sites were obtained across all patients, noting that for the most superficial recording sites within the STN, the macroelectrode ring may have been just superior to the dorsal border of STN. The locations of the macroelectrode contacts were determined using the semi-automatic approach implemented in the Lead-DBS toolbox (
Horn and Kühn, 2015; Horn et al., 2019). In brief, post-operative CT scans were linearly coregistered with pre-operative MRI scans and normalized to MNI (Montreal Neurological Institute) space. MNI-defined coordinates of macroelectrode contact locations were extracted for all subjects and visualized in Figure 2B.

**Figure 2.**
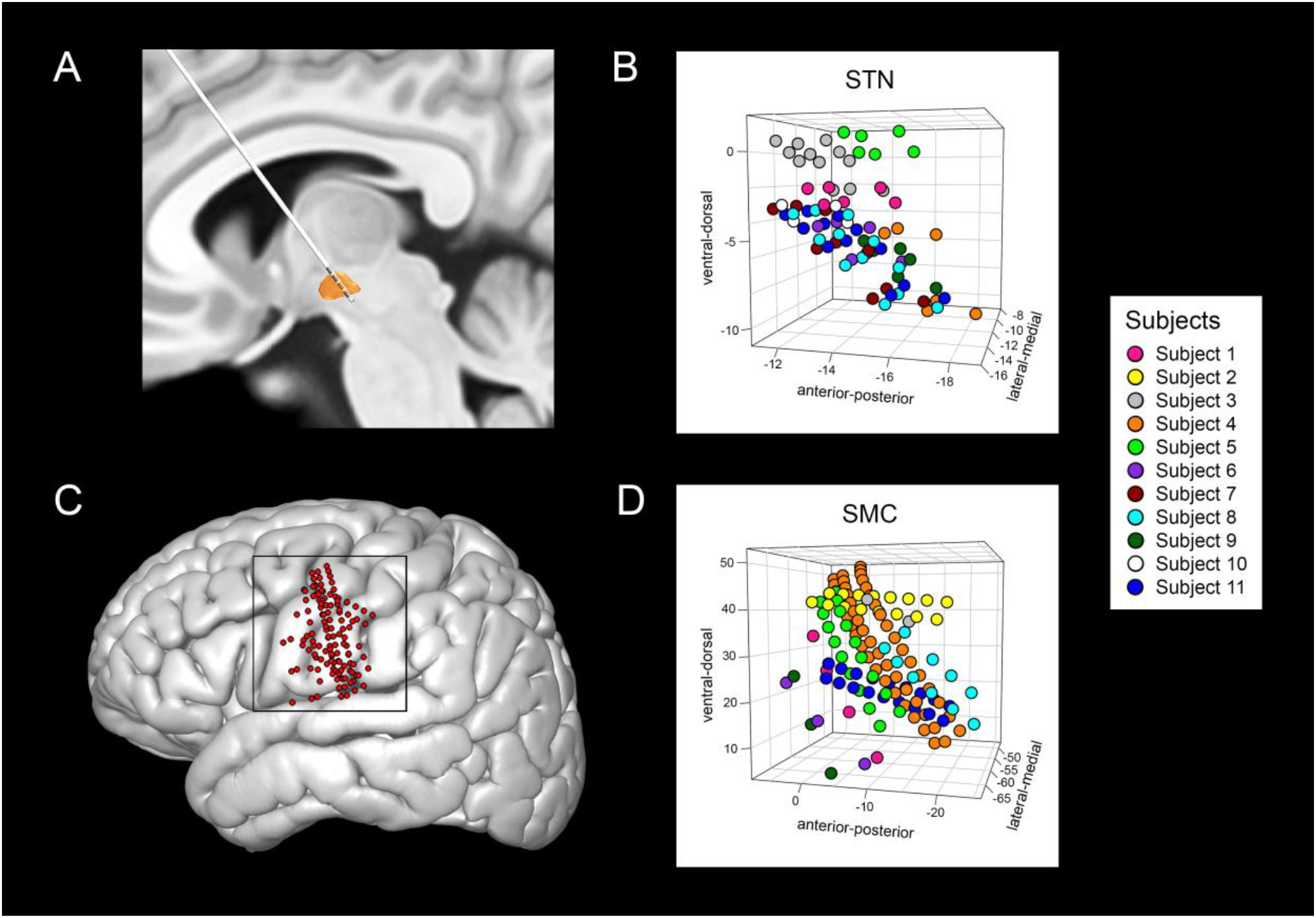
Location of recording sites in the MNI-defined space. ***A***, An example trajectory of the DBS lead through the left subthalamic nucleus (STN) shown on the DISTAL atlas by Ewert et al. (2017). ***B***, MNI-defined coordinates (mm) of recording sites in the STN plotted for all subjects in 3D space. ***C***, Reconstructed locations of all ECoG electrodes in the sensorimotor cortex that were included in the study (n = 125), co-registered and plotted on the cortical surface of the MNI brain space. ***D***, MNI-defined coordinates (mm) of the ECoG contacts on the sensorimotor cortex plotted for all subjects in 3D space. In ***B*** and ***D***, each subject’s electrodes are mapped with a different color.

### Cortical recordings

In addition to the clinical subcortical mapping procedure, all patients were also temporarily implanted with subdural electrode arrays over the cortical surface of the left hemisphere which were inserted through the burr hole after opening the dura, but before the insertion of subcortical guide tubes. The ECoG signal was acquired at 30 kHz using the Grapevine Neural Interface Processor. Most subjects were implanted with 6- or 28-channel Ad-Tech electrode strips (Ad-Tech Medical Corporation, Racine, WI, USA) except for two subjects who were implanted with either a 36- or 54-channel PMT electrode strips each (PMT Corporation, Chanhassen, MN, USA). Depending on the type of the electrodes, the electrodes varied 1, 2 or 4 mm in diameter, and 3, 4 or 10 mm in center-to-center spacing. The placement of the electrode strips was targeted at the ventral sensorimotor cortex by using stereotactic coordinates to mark the scalp over this region and advancing the subdural strips in the direction of this overlying visual marker. A total of 198 electrodes were placed on the cortex, but only 125 were included in the analyses – those that were confined to the sensorimotor cortex, as determined in the patients’ native brain space (Figure 2C shows these locations in MNI space). Localization of the electrodes on the cortical surface was reconstructed from 1) the intra-operative fluoroscopic images (512 × 512 pixels, General Electric, OEC 9900) and 2) the coregistered pre-operative and post-operative computed tomography (CT) images obtained after placement of the Leksell frame and 3) pre-operative magnetic resonance imaging (MRI) scans according to the semi-automated method described in Randazzo et al. (2016). Electrode locations were then registered to the common brain space using the MNI template (ICBM152) with Brainstorm (Tadel et al., 2011) (https://neuroimage.usc.edu/brainstorm/). Subjects’ MNI-defined ECoG electrodes that were constrained to the sensorimotor cortex in native space are presented in 3D MNI space in Figure 2D.

### Data selection

Of the 11 subjects who participated in the study, STN data for one subject (Subject 2) was not recorded due to a technical error. ECoG data from 2 subjects contained excessive artifacts in the signal (Subjects 7 and 10) and were excluded from the analysis. Trials were included in the analysis if 1) a student coding the data was able to unambiguously identify a subject’s spoken response; 2) a subject’s spoken response constituted the stimuli’s targeted CVC structure; 3) a subject’s response included the stimuli’s targeted phonemes. On the basis of these criteria, 359 (9.8%) out of a total of 3,669 recorded trials were rejected.

### Electrophysiological data processing

Data processing was performed using custom code based on the Statistical Parametric Mapping (SPM12) (Wellcome Department of Cognitive Neurology, London, UK) (http://www.fil.ion.ucl.ac.uk/spm/software/spm12/) and Fieldtrip (Oostenveld et al., 2011) toolboxes implemented in MATLAB. The data were resampled to a sampling frequency of 1 kHz. In order to minimize noise and artifactual electrode cross-talk in the signal, the data were re-referenced offline using a common average referencing procedure applied over blocks of electrodes connected by the same headstage connector for the ECoG recordings, and using a common average referencing procedure for the STN recordings. A 1 Hz high-pass filter and a 58-62 Hz notch filter were applied to remove cardioballistic artifacts and line noise, respectively. The signal was then aligned with the presentation of the green cross cue for subsequent baseline epoching and with the vowel onset (the transition between the initial consonant and the subsequent vowel, CV) for speech response epoching. The CV transition was used to separate the consonantal component from the subsequent vocalic component in subjects’ spoken responses (as in Bouchard et al., 2013). For artifact rejection, data were visually inspected over 6000-ms long time windows surrounding baseline and vowel onset; time widows with residual artifacts and excessive noise were excluded from analysis, resulting in an additional 4.8% data rejection. The remaining data underwent a time-frequency transformation using Morlet wavelets with 7 cycles over frequencies between 1 and 200 Hz in incrementing steps of 2 Hz. The resulting signal was normalized using z-scores calculated relative to a 1000-ms long baseline period (250 ms before and 750 ms after green cross presentation). A time-varying analytic amplitude in the high gamma frequency range (60-150 Hz) was extracted for further analyses because it has been consistently reported to reflect changes in sensory, motor and cognitive functions, including speech (e.g., Bouchard et al., 2013; Crone et al., 1998; Edwards et al., 2005).

### Experimental design and statistical analysis

All statistical analyses were performed in MATLAB 2017a and R version 3.4.4 (R Development Core Team, 2018). A within-subjects experimental design was used, in which all subjects (n = 11) received trials with both lips and tongue articulations. Recorded LFPs from the sensorimotor cortex and the subthalamic nucleus were analyzed separately using the same statistical procedures. For the analysis of the LFPs throughout speech production, a time window of 1000 ms (500 ms before and 500 ms after the vowel onset) encompassing subjects’ whole spoken response was used. For the analysis of the articulatory specificity of the initial consonant, a 500-ms time window preceding the vowel onset was used. Although the duration of the word-initial consonant in the prevocalic position can vary in the range of 20-150 ms depending on the type of consonant and subsequent vowel (Umeda, 1977; Son and Santen, 1997), a broader time window of 500 ms allows examination of potential pre-articulatory neuronal activity. To analyze high gamma activity elicited during the speech task, a series of fitted linear mixed effects models (LMEMs) with restricted maximum likelihood estimation were carried out using lme4 (Bates, Maechler, Bolker, and Walker, 2015) and lmerTest (Kuznetsova, Brockhoff, and Christensen, 2017) packages. Subjects were entered as random effects to account for subject-specific idiosyncrasies. Model comparisons were performed via backward elimination of fixed effects and their interactions in order to measure the goodness of model fit without unnecessary parameter overfitting using the Akaike information criterion (Akaike, 1974). To perform correlation analyses between the observed speech and articulatory response and electrode location coordinates in the MNI space, we applied a Spearman’s rank correlation test. Generally, to assess statistical differences of speech-related changes in the brain response, we used Welch two sample t-tests when the data were found not to deviate significantly from normality (as determined by a Shapiro-Wilk normality test); when the data were not normally distributed, nonparametric Wilcoxon rank sum or Wilcoxon signed-rank (to determine the significance of response compared to baseline) tests were used. False discovery rate (FDR) method (as described in Benjamini and Hochberg, 1995) was used at *α* = 0.05 to control for multiple comparisons.

## RESULTS

### Behavioral response

Subjects’ behavioral performance is summarized in Table 2. Across subjects, the mean latency from seeing the stimulus word on the screen to producing the word was 1.34 ± 0.51 s; the mean duration of the spoken response was 0.59 ± 0.16 s. The severity of the disease symptoms as measured by the UPDRS off medication did not account for variation in response latency (estimated coefficient = −0.006, SE = 0.018, *t* = −0.23, *p* = 0.77) or response duration (estimated coefficient = −0.003, SE = 0.005, *t* = −0.63, *p* = 0.54). Average response accuracy was 88.5%, although subjects 1, 2, and 7 produced many non-target responses (incomplete words and/or non-target phonemes) resulting in a high percent of rejected trials (more than 20%).

**Table 2.**
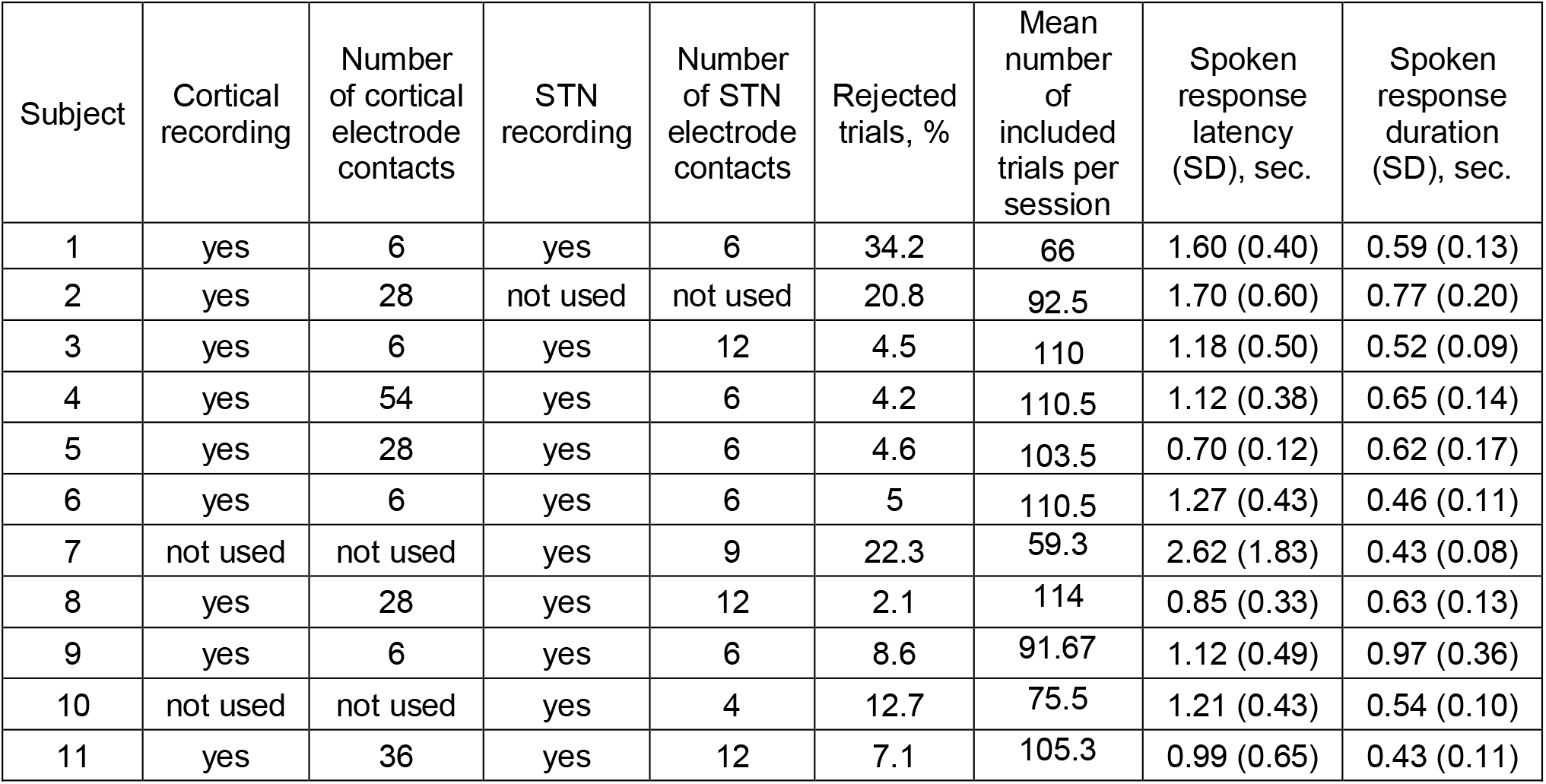
Subjects recording and behavioral performance characteristics.

| Subject | Cortical recording | Number of cortical electrode contacts | STN recording | Number of STN electrode contacts | Rejected trials, % | Mean number of included trials per session | Spoken response latency (SD), sec. | Spoken response duration (SD), sec. |
| --- | --- | --- | --- | --- | --- | --- | --- | --- |
| 1 | yes | 6 | yes | 6 | 34.2 | 66 | 1.60 (0.40) | 0.59 (0.13) |
| 2 | yes | 28 | not used | not used | 20.8 | 92.5 | 1.70 (0.60) | 0.77 (0.20) |
| 3 | yes | 6 | yes | 12 | 4.5 | 110 | 1.18 (0.50) | 0.52 (0.09) |
| 4 | yes | 54 | yes | 6 | 4.2 | 110.5 | 1.12 (0.38) | 0.65 (0.14) |
| 5 | yes | 28 | yes | 6 | 4.6 | 103.5 | 0.70 (0.12) | 0.62 (0.17) |
| 6 | yes | 6 | yes | 6 | 5 | 110.5 | 1.27 (0.43) | 0.46 (0.11) |
| 7 | not used | not used | yes | 9 | 22.3 | 59.3 | 2.62 (1.83) | 0.43 (0.08) |
| 8 | yes | 28 | yes | 12 | 2.1 | 114 | 0.85 (0.33) | 0.63 (0.13) |
| 9 | yes | 6 | yes | 6 | 8.6 | 91.67 | 1.12 (0.49) | 0.97 (0.36) |
| 10 | not used | not used | yes | 4 | 12.7 | 75.5 | 1.21 (0.43) | 0.54 (0.10) |
| 11 | yes | 36 | yes | 12 | 7.1 | 105.3 | 0.99 (0.65) | 0.43 (0.11) |

### Speech-related activity

STN LFP activity showed significant time-frequency modulations relative to baseline (Figure 3A) that were comparable with those obtained from the sensorimotor cortex (Figure 3B). There were significant decreases in z-scored spectral power in the alpha (8-12 Hz) and beta (12-30 Hz) frequency bands and significant increases in z-scored power at high frequency ranges relative to baseline, as determined by the Wilcoxon signed-rank test (*α* = 0.05, FDR corrected). Increases in the spectral power occurred from 60 Hz to180 Hz for STN sites, and from 50Hz and onward for the cortical sites. In both cases, significant high frequency modulations occurred around 400 ms before speech onset and persisted until about 100 ms before speech offset for STN activity and until about 100 ms after speech offset for sensorimotor cortex activity. A more detailed examination of the z-scored spectral power in the high gamma frequency range over the spoken response window (500 ms before vowel onset and 500 ms after vowel onset) showed that in 86% (68/79) of STN sites and 95% (119/125) of sensorimotor cortex ECoG sites, high gamma power was significantly greater than baseline (Wilcoxon signed-rank test at *α* = 0.05, FDR corrected). The subjects’ symptom severity (as measured by a total UPDRS score) was not correlated (Spearman’s rank-order correlation test) with the average high gamma activity for the speech response window either in the STN (*r_s_*(10) = 0.37, *p* = 0.29) or the sensorimotor cortex (*r_s_*(9) = 0.43, *p* = 0.24). In the STN, averaged high gamma power significantly correlated with the location of recording sites along the ventral-dorsal axis in the MNI space (*r_s_*(79) = 0.53, *p* < 0.001) and anterior-posterior axis (*r_s_*(79) = 0.36, *p* = 0.0012), but not the lateral-medial axis (*r_s_*(79) = 0.03, *p* = 0.78). In contrast, we found no significant correlation between high gamma power and the location of the recording sites on the sensorimotor cortex. To explain the observed variation in the high gamma power across STN recording sites and to control for subject variability, we fitted linear mixed effects models (LMEM). The most parsimonious model included average high gamma power as a dependent variable, subjects as a random effect, and the location of recording sites along the ventral-dorsal axis (the MNI-defined Z coordinate) as a fixed effect. The outcome of the LMEM suggests that, even after taking subject-to-subject variability into account, high gamma power changed significantly from dorsal to ventral parts of the STN, with greater high gamma power observed dorsally (estimated coefficient = 0.017, SE = 0.005, *t* = 3.19, *p* = 0.004). Mixed effects modeling of the high gamma response in the sensorimotor cortex did not yield significant effects.

**Figure 3.**
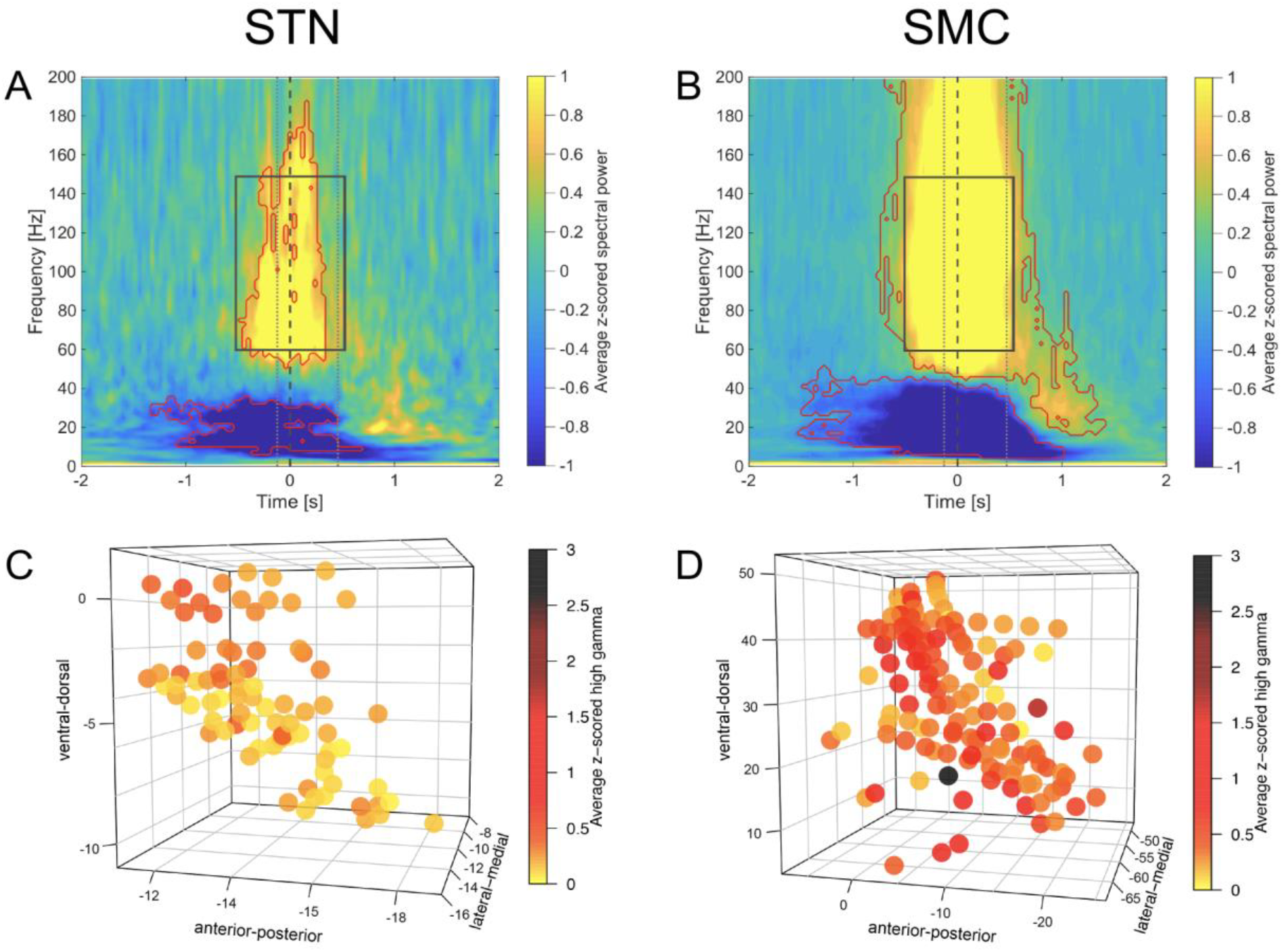
Subthalamic nucleus (STN) and sensorimotor cortex (SMC) show speech production-related time-frequency modulations. ***A-B***, Grand average of (***A***) STN and (***B***) SMC oscillatory activity (average z-scored spectral power) across all recording sites and all trials aligned to vowel onset (Time = 0 s, grey dashed vertical line). Significant modulations compared to baseline are marked in red contour (Wilcoxon signed-rank, *p* < 0.05, FDR corrected). Average speech production onsets and offsets are marked with grey dotted vertical lines. Rectangles with grey solid lines mark the time window (±500 ms from vowel onset) for the analysis of speech production-related high gamma (60-150 Hz) activity. ***C-D***, Z-scored high gamma (60-150 Hz) power averaged for the 1 s time window (±500 ms from vowel onset) plotted in 3D space for each subject’s (***C***) STN and (***D***) SMC recording site. The location of recoding sites is provided in MNI coordinates.

### Representation of articulators

To examine the spatial distribution of speech articulator representations within the STN and sensorimotor cortex, we compared (Welch two sample t-test) high gamma power in trials with the tongue as the main articulator vs. trials with the lips as the main articulator, at each recording site. We used the outcome of the t-test and the sign of the t-value to detect discriminative articulatory activity. For example, a significant (*α* = 0.05) and positive t-value indicated that a given site’s high gamma power was greater for consonants articulated with the lips than those articulated with the tongue; conversely, negative t-values indicated tongue-related activity. Eighteen STN sites (23%) out of a total of 79 were discriminative: 5 sites exhibited greater high gamma activity during articulation with the lips; 13 sites were most active during articulation with the tongue (Figure 4A). Thirty-seven sites (30%) out of a total of 125 sites on the sensorimotor cortex showed lips-preferred activity (n = 19) or tongue–preferred activity (n = 18) (Figure 4C). An example of what constituted an articulator-preferred activity is shown for representative recording sites in Figures 4B and D. The remaining sites at which a significant increase in high gamma power was observed produced an undifferentiated activity, i.e. they were equally active during articulation of both coronal and labial consonants. In the STN, recording sites with a tongue-preferred response appeared potentially to be located more dorsally compared to those selective for lips; however, the obtained t-values did not correlate significantly with any of three spatial orientation planes through the recording locations (ventrodorsally, anteroposteriorly, or lateromedially), according to a Spearman’s rank-order correlation test. Modeling of the articulatory activity in the STN with mixed effects regression approach did not yield significant effects (the most parsimonious model included subjects as a random effect and recording locations along the lateral-medial axis (the MNI-defined X coordinate) as a fixed effect). In the sensorimotor cortex, t-values correlated significantly with the location of recording sites along the ventral-dorsal (*r_s_*(125) = −0.39, *p* < −0.001) and lateral-medial (*r_s_*(125) = −0.35, *p* = < 0.001) axes. Modeling the articulatory effect with LMEMs produced similar results. Keeping subjects as a random effect, the most parsimonious models yielded a significant effect of the recording location along the ventral-dorsal (estimated coefficient = 0.064, SE = 0.022, *t* = 2.98, *p* = 0.004) and the lateral-medial (estimated coefficient = 0.148, SE = 0.053, *t* = 2.6, *p* = 0.011) axes. Thus, taking subject-to-subject differences into account, the articulator-related activity in the sensorimotor cortex appeared to be somatotopically organized, with the recording sites exhibiting encoding of lip articulations located more dorsally (and medially due to the cortex curvature), and sites exhibiting encoding of tongue articulations distributed more broadly over the ventrolateral part of the sensorimotor cortex.

**Figure 4.**
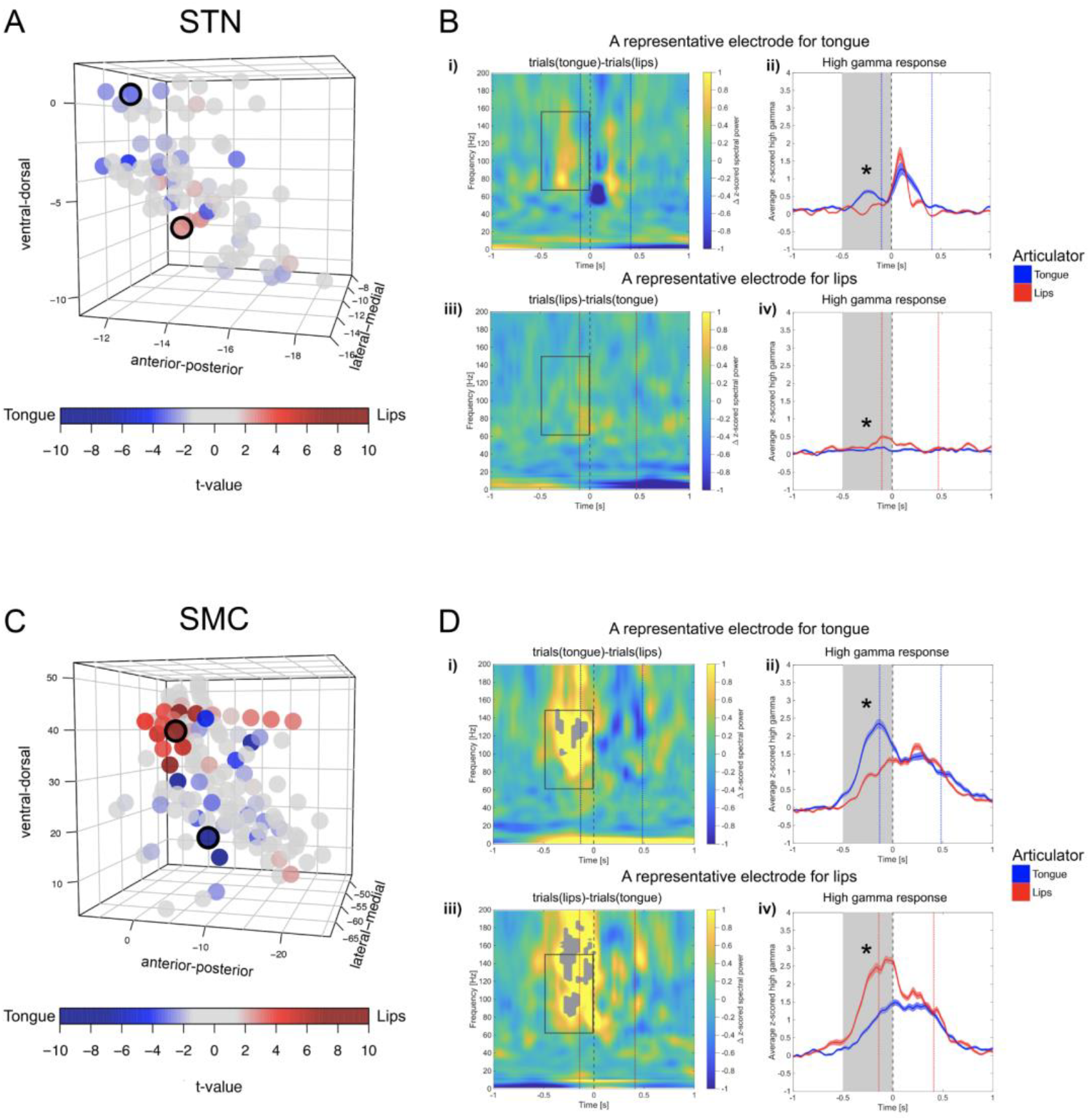
Spatial distribution of tongue- and lips-preferred articulatory activity in the MNI-defined STN space and sensorimotor cortex (SMC). ***A*** and ***C***, Outcome of a series of t-tests comparing z-scored high gamma power (averaged for a 0.5 s long time window before vowel onset) during articulation of tongue consonants vs. lips consonants for each (***A***) STN and (***C***) SMC recording site. Opacity of the circles varies with the magnitude of the t-value: negative t-values (in blue shades) suggest a greater response to tongue; positive t-values (in red shades) suggest a greater response to lips (Welch two sample t-test, *p* < 0.05). Note that the obtained t-values for the SMC sites differed significantly along the ventral-dorsal and lateral-medial axes (Spearman’s rank-order correlation test, *p*< 0.01), suggesting articulator-discriminative somatotopy. Circles with black outline mark representative sites for tongue and lips, whose articulatory activity is plotted on the right. ***B*** and ***D***, Examples of representative tongue-preferred and lips-preferred sites for (***B***) STN and (***D***) SMC. A subtraction time-frequency representation is shown for (***i***) the tongue-preferred site after time-frequency representation for all trials with lips consonants is subtracted from time-frequency representation for all trials with tongue consonants, and for (***iii***) the lips-preferred site after time-frequency representation for all trials with tongue consonants is subtracted from time-frequency representation for all trials with lips consonants. Grey filled contours mark significant time-frequency differences between the two conditions (Wilcoxon rank sum test, *p* < 0.05, FDR corrected). Rectangles with grey solid lines mark the time window (from 0.5 s before vowel onset until vowel onset) for the analysis of articulator-specific high gamma (60-150 Hz) activity. Differences in averaged z-scored high gamma power elicited by trials with the tongue articulation vs. the lips articulation are shown for tongue-specific (***ii***) and lips-specific (***iv***) sites (significant differences are marked with asterisks, Welch two sample t-test, *p* < 0.05). Gray bands mark the time window (from 0.5 s before vowel onset until vowel onset) across which high gamma power was averaged for the analysis of articulator-specific activity. Throughout ***i-iv***, grey dashed vertical line represents vowel onset (Time = 0 s). Dotted vertical lines represent spoken response onsets and offsets for trials with tongue consonants (blue) and trials with lips consonants (red).

To examine the time-course of articulatory neural encoding, we first compared amplitude and time of peak high gamma activity in the identified articulator-discriminative sites in the sensorimotor cortex and the STN. Because there was no significant difference in the average or peak high gamma amplitude between tongue vs. lips-preferred sites, high gamma activity was plotted for combined (tongue and lips) articulator-discriminative sites (Figure 5A). Mean amplitude of high gamma activity was significantly different between cortical (Mean = 0.36, SD = 0.49) and STN (Mean = 0.16, SD = 0.15) recording sites (*t*(2478) = 16.66, *p* < 0.001), but the times of peak high gamma activity did not differ between the two structures (sensorimotor cortex (Mean = 0.017 s, SD = 0.22 s) and STN (Mean = 0.004 s, SD = 0.21 s), *t*(35.65) = 0.21, *p* = 0.84). Next, for each articulator-discriminative recording site, we carried out a Welch two sample t-test for the high gamma power in tongue vs. lips trials at each time point (n = 51, Δt = 40 ms) within the 2 s interval centered at vowel onset. Significant t-test outcomes (*p* < 0.05) indicating presence of articulatory discrimination at a given time point are plotted in Figure 5B. Such analytic strategy enabled interpretation of articulatory encoding regardless of the underlying high gamma amplitude. For example, negative and positive t-values with *p* < 0.05 indicated tongue-preferred or lip-preferred activity, while t-values close to zero indicated indiscriminate articulatory activity because the underlying high gamma spectral power for labial and coronal consonants was similar (regardless of its amplitude). We found that the timing of the articulatory encoding varied across responsive electrodes, but was, overall, less variable in the sensorimotor cortex compared to the STN. In the sensorimotor cortex, high gamma amplitude in tongue vs. lips trials started to diverge significantly well before articulations (consistent with Bouchard et al. (2013)), however the bulk of significant observations concentrated around the time of consonant onset (Figure 5B). In the STN, significant articulator-discriminative observations were more broadly spread, often having two peaks. The number of obtained significant outcomes for each time point of the response was significantly greater for the cortical (Mean = 6.43, SD = 6.9) than the STN (Mean = 2.85, SD = 1.7) recording sites (*t*(51.7) = 3.43, *p* = 0.0012), but the times at which articulatory encoding (significant t-values) was observed on average did not differ between the two structures (sensorimotor cortex (Mean = −0.052 s, SD = 0.53 s) and STN (Mean = 0.015 s, SD = 0.38 s), *t*(189.4) = −1.29, *p* = 0.2).

**Figure 5.**
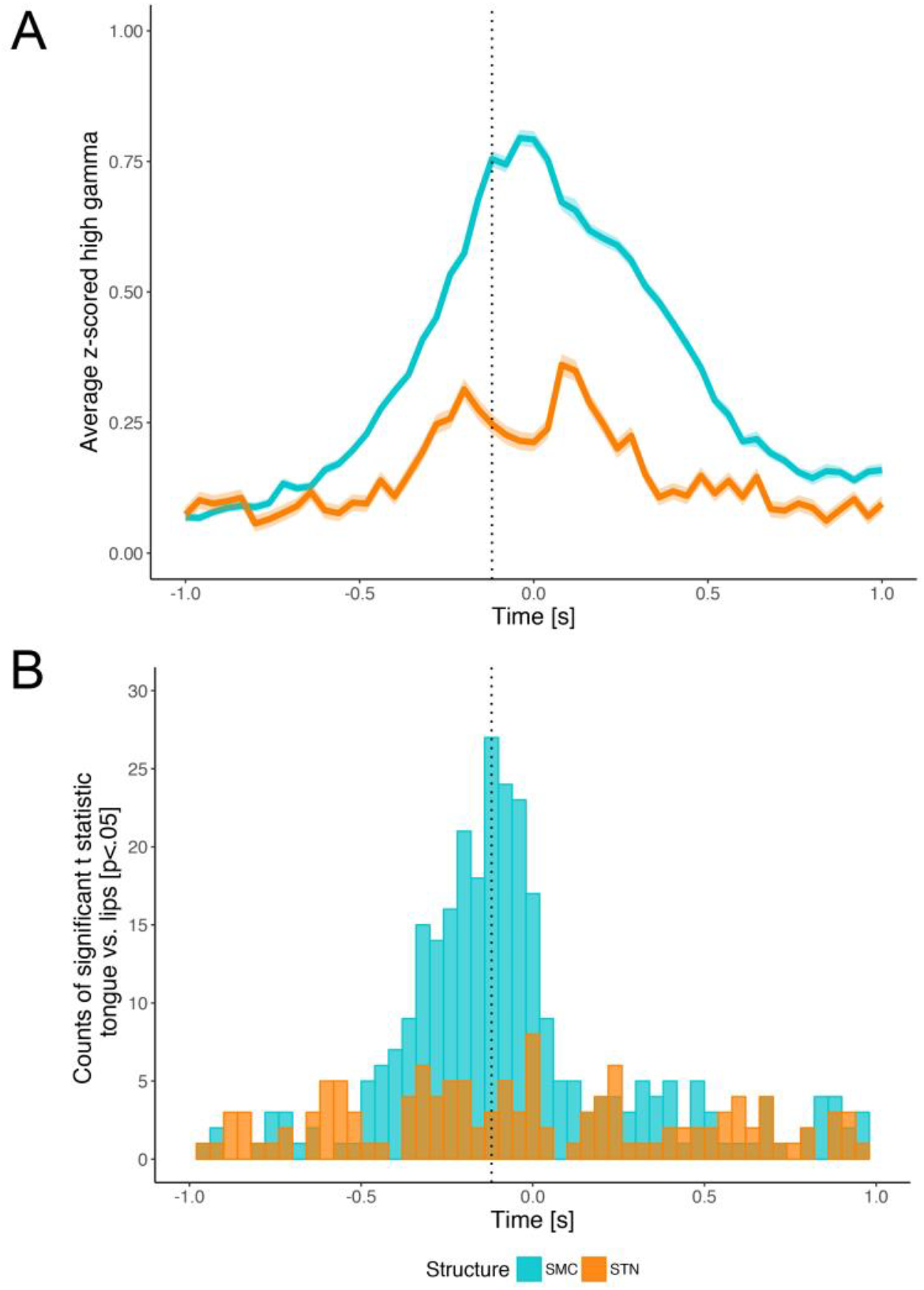
Time-course of the articulatory encoding in the subthalamic nucleus (STN) and sensorimotor cortex (SMC). ***A***, Average high gamma activity in the STN and SMC articulator-discriminative recording sites. ***B***, Distribution of significant outcomes (*p* < 0.05) of the Welch two sample t-test comparing z-scored high gamma power during articulation of tongue consonants vs. lips consonants at each time point of the STN and the SMC recordings. ***A-B***, Dotted vertical lines represent consonant onset; vowel onset is at Time = 0 s.

## Discussion

We analyzed local field potentials obtained from the simultaneous recording of the cortical and STN activity in 11 patients with Parkinson’s disease while they participated in a speech task during subcortical mapping for the implantation of DBS electrodes. We selected the speech stimuli such that articulation of the initial consonant engaged either tongue or lip musculature, in order to examine whether encoding of speech articulators, similar to that previously reported for the sensorimotor cortex, is represented in subthalamic high gamma activity. We found that STN high gamma activity does track speech production at the level of vocal tract articulators, with similar temporal, but different spatial organization of articulatory neural representations compared to that of the sensorimotor cortex.

### Speech-related activation

We found that speech production was accompanied by significant time-frequency modulations in both the STN and the sensorimotor cortex, namely, suppression of alpha and beta activity and increase in high gamma activity (above 50 Hz). In both cases, significant time-frequency modulations emerged about 400 ms before spoken response onset and persisted throughout the execution of speech. Desynchronization of alpha and beta activity and synchronization of high frequency activity have been previously reported as markers of ongoing movement and movement-related patterns in the STN (Androulidakis et al., 2007; Kempf et al., 2007; Lipski et al., 2017; Geng et al., 2018; Lofredi et al., 2018). However, only modulation of beta activity during speech production has been reported (Hebb, Darvas, and Miller, 2012). Thus, our results provide the first demonstration of high gamma synchronization before and during speech production in STN. Importantly, we show that the power of high gamma response changes significantly along the dorsoventral plane of the MNI-defined locations of the STN electrodes, with greater high gamma power observed dorsally. This finding agrees with recent demonstration that subthalamic gamma power is greatest in the sensorimotor part of the subthalamic nucleus (Lofredi et al., 2018). Thus, in light of the existing conception of the parcellated organization of the STN into sensorimotor, associative, and limbic areas (Hamani et al., 2004; Temel et al., 2005; Haynes and Haber, 2013), our results show that articulatory aspects of speech recruit the sensorimotor region of the STN, and are in line with our previous findings showing speech-related increases in the firing rate of human STN neurons (Lipski et al., 2018). In contrast to the STN, the magnitude of cortical high gamma activity was not significantly different across recording locations. Given that the cortical recordings were confined to the orofacial segment of the sensorimotor cortex and the evidence of overlapping speech-related activation in the precentral and postcentral gyri (Cheung et al., 2016; Conant et al., 2018), this lack of spatial differentiation in the cortical high gamma activity is expected.

### Encoding of speech articulators

In order to further quantify the observed speech-related high gamma modulation in the STN and the sensorimotor cortex, we examined whether the two structures showed encoding specific to speech articulators. For the sensorimotor cortex, we found that 30% of recording sites revealed either lips-preferred or tongue-preferred activity, which had a topographic distribution: the electrodes located more dorsally on the sensorimotor cortex produced a greater high gamma power for the articulation of lips consonants while the electrodes that were located more ventrally yielded a greater high gamma power for tongue consonants. Thus, our results appear to recapitulate the expected dorsoventral layout for lips and tongue representations within the primary motor and sensory cortices (Penfield and Boldrey, 1937; Bouchard et al., 2013; Breshears et al., 2015; Chartier et al., 2018; Conant et al., 2018). We found that the density of articulator-discriminative observations closely aligned with the consonant onset in acoustic speech production. This discriminative activity began to emerge about 500 ms before articulation, suggesting the potential encoding of pre-articulatory preparatory processes like planning a motor command and retrieving the sensory representation of the intended articulatory target (Guenther, Ghosh, and Tourville, 2006). For the subthalamic nucleus, we found that 23% of recording locations showed articulator-discriminative activity, with temporal onset also occurring before articulation, but without articulatory somatotopy. Previous studies demonstrating functional organization in the STN of human and nonhuman primates have used single unit recordings to demonstrate a crude somatotopy for arm-related and leg-related movements (Monakow et al., 1978; DeLong et al., 1985; Wichmann et al., 1994; Nambu, Takada, Inase, and Tokuno, 1996; Rodriguez-Oroz et al., 2001; Starr et al., 2003; Theodosopoulos et al., 2003), although reported representations for face, eyes and finer-grained movements with shoulders, elbows, knees, wrists, etc. have been less somatotopically consistent (e.g., DeLong et al., 1985; Wichmann et al., 1994). It should be noted that LFP recordings might not be expected to delineate a functional somatotopy, due to their representation of group level neuronal activity recorded from a much larger volume of tissue than the signal obtained from microelectrode recordings. In this respect, it is remarkable that we found evidence for articulator-level encoding in the LFP signal, which may indicate the encoding of aspects of speech production that are specific to these articulatory maneuvers but separate from their anatomical representation at the cortical level, such as effort or switching between motor plans (Redgrave, Prescott, and Gurney, 1999; Frank, 2006). Our study remains agnostic as to the exact computational processes underlying the observed articulator-related differences: it remains to be established whether they reflect musculature engaged in articulation, motor planning, control over kinematic trajectories, or alternatively higher-order speech-related processes.

#### Limitations

We acknowledge that the disease state is a potential confound to our results. We do not report control data collected from a non-PD population. Given that basal ganglia activity in PD patients is characterized by reorganization of receptive fields and loss of specificity (Abosch et al., 2002; Hamani et al., 2004), we may be assessing an unknown amount of crosstalk or “motor overflow” of the signal related to different body parts (Bergman et al., 1998; Nambu, 2011). Additionally, we searched for articulator-specific somatotopy on the basis of 79 available STN recording locations with non-systematic spatial separation, which may represent inadequate sampling. Note that fine-wire EMG is not an option in awake neurosurgical patients, thus our experimental design did not allow us to measure articulatory muscle movement for correlation with intracranial signals.

#### Summary

These data are the first to demonstrate time-frequency modulations in STN activity that tract articulatory aspects of speech, corroborating recent evidence for speech production-related changes in the timing and the firing rate of the STN neurons (Lipski et al., 2018). A major strength of this study is that, by applying the same methodological and analytical approach to analyze simultaneous LFP recordings from the sensorimotor cortex and the STN, we could compare the neural response in the STN to that obtained from the sensorimotor cortex. After demonstrating the expected somatotopic differentiation of vocal tract articulators in the sensorimotor cortex, we showed that the STN also encodes speech articulators. Further elucidation of the role of cortico-basal ganglia interactions in the speech production network will be critical for improving our understanding of the neurobiology of speech dysfunction in basal ganglia disorders and related future treatments.

## Acknowledgements

Funding was provided by NINDS U01NS098969 (PI: Richardson), the Hamot Health Foundation (PI: Richardson), and a University of Pittsburgh Brain Institute NeuroDiscovery Pilot Research Award (PI: Richardson). The authors are thankful to the patients who participated in the study, the clinical and the operating room staff for help with data collection.

## REFERENCES

Abosch A, Hutchison WD, Saint-Cyr JA, Dostrovsky JO, Lozano AM (2002) Movement-related neurons of the subthalamic nucleus in patients with Parkinson disease. J Neurosurg 97:1167–1172.

Akaike H (1974) A new look at the statistical model identification. IEEE Transactions on Automatic Control 19:716–723.

Aldridge D, Theodoros D, Angwin A, Vogel AP (2016) Speech outcomes in Parkinson’s disease after subthalamic nucleus deep brain stimulation: A systematic review. Parkinsonism Relat Disord 33:3–11.

Alexander GE, DeLong MR, Strick PL. (1986) Parallel organization of functionally segregated circuits linking basal ganglia and cortex. Annu Rev Neurosci 9:357–81.

Androulidakis AG, Kühn AA, Chen CC, Blomstedt P, Kempf F, Kupsch A, … Brown P (2007) Dopaminergic therapy promotes lateralized motor activity in the subthalamic area in Parkinson’s disease. Brain, 130:457–468.

Bates D, Maechler M, Bolker B, Walker S (2015) Fitting linear mixed-effects models using lme4. Journal of Statistical Software 67:1–48.

Benjamini Y, Hochberg Y (1995) Controlling the false discovery rate: A practical and powerful approach to multiple testing. Journal of the Royal Statistical Society 57:289–300.

Bergman H, Feingold A, Nini A, Raz A, Slovin H, Abeles M, Vaadia E (1998) Physiological aspects of information processing in the basal ganglia of normal and parkinsonian primates. Trends in neurosciences 21:32–38.

Bouchard KE, Chang EF (2014) Control of spoken vowel acoustics and the influence of phonetic context in human speech sensorimotor cortex. Journal of Neuroscience 34:12662–12677.

Bouchard KE, Conant DF, Anumanchipalli GK, Dichter B, Chaisanguanthum KS, Johnson K, Chang EF (2016) High-resolution, non-invasive imaging of upper vocal tract articulators compatible with human brain recordings. PLoS One 11:e0151327.

Bouchard KE, Mesgarani N, Johnson K, Chang EF (2013) Functional organization of human sensorimotor cortex for speech articulation. Nature 495:327–332.

Brainard DH (1997) The Psychophysics Toolbox. Spatial Vision 10:433–436.

Breshears JD, Molinaro AM, Chang EF (2015) A probabilistic map of the human ventral sensorimotor cortex using electrical stimulation. Journal of Neurosurgery 123:340–349.

Brown S, Ngan E, Liotti M (2007) A larynx area in the human motor cortex. Cerebral Cortex 18:837–845.

Brunner RJ, Kornhuber HH, Seemtiller E, Suger G, Wallesch CW (1982) Basal ganglia participation in language pathology. Brain and Language 16:281–299.

Carey D, Krishnan S, Callaghan MF, Sereno MI, Dick F (2017) Functional and quantitative MRI mapping of somatomotor representations of human supralaryngeal vocal tract. Cerebral Cortex 27:265–278.

Chartier J, Anumanchipalli GK, Johnson K, Chang EF (2018) Encoding of articulatory kinematic trajectories in human speech sensorimotor cortex. Neuron 98:1042–1054.

Cheung C, Hamilton LS, Johnson K, Chang EF (2016) The auditory representation of speech sounds in human motor cortex. Elife 5:e12577.

Conant DF, Bouchard KE, Leonard MK, Chang EF (2018) Human sensorimotor cortex control of directly-measured vocal tract movements during vowel production. Journal of Neuroscience 2382–17.

Crone NE, Miglioretti DL, Gordon B, Lesser RP (1998) Functional mapping of human sensorimotor cortex with electrocorticographic spectral analysis: II. Event-related synchronization in the gamma band. Brain 121:2301–2315.

Damasio AR, Damasio H, Rizzo M, Varney N, Gersh F (1982) Aphasia with nonhemorrhagic lesions in the basal ganglia and internal capsule. Arch Neurol 39:15–24.

DeLong MR, Crutcher MD, Georgopoulos AP (1985) Primate globus pallidus and subthalamic nucleus: functional organization. Journal of Neurophysiology 53:530–543.

Edwards E, Soltani M, Deouell LY, Berger MS, Knight RT (2005) High gamma activity in response to deviant auditory stimuli recorded directly from human cortex. Journal of Neurophysiology 94:4269–4280.

Ewert S, Plettig P, Li N, Chakravarty MM, Collins DL, Herrington TM, … Horn A (2017) Toward defining deep brain stimulation targets in MNI space: A subcortical atlas based on multimodal MRI, histology and structural connectivity. NeuroImage 170: 271–282.

Frank MJ (2006) Hold your horses: A dynamic computational role for the subthalamic nucleus in decision making. Neural Networks 19:1120–1136.

Geng X, Xu X, Horn A, Li N, Ling Z, Brown P, Wang S (2018) Intra-operative characterisation of subthalamic oscillations in Parkinson’s disease. Clinical Neurophysiology 129:1001–1010.

Guenther FH, Ghosh SS, Tourville JA (2006) Neural modeling and imaging of the cortical interactions underlying syllable production. Brain and Language 96:280–301.

Hamani C, Saint-Cyr JA, Fraser J, Kaplitt M, Lozano AM (2004) The subthalamic nucleus in the context of movement disorders. Brain 127:4–20.

Haynes WI, Haber SN (2013) The organization of prefrontal-subthalamic inputs in primates provides an anatomical substrate for both functional specificity and integration: implications for Basal Ganglia models and deep brain stimulation. Journal of Neuroscience 33:4804–4814.

Hebb AO, Darvas F, Miller KJ (2012) Transient and state modulation of beta power in human subthalamic nucleus during speech production and finger movement. Neuroscience 202:218–233.

Hesselmann V, Sorger B, Lasek K, Guntinas-Lichius O, Krug B, Sturm V, … Lackner K (2004) Discriminating the cortical representation sites of tongue and lip movement by functional MRI. Brain Topography 16:159–167.

Ho AK, Iansek R, Marigliani C, Bradshaw JL, Gates S (1998) Speech impairment in a large sample of patients with Parkinson’s disease. Behav Neurol 11:131–137.

Horn A, Kühn AA (2015) Lead-DBS: a toolbox for deep brain stimulation electrode localizations and visualizations. Neuroimage 107:127–135.

Horn A, Li N, Dembek TA, Kappel A, Boulay C, Ewert S, … Reisert M (2019) Lead-DBS v2: Toward a comprehensive pipeline for deep brain stimulation imaging. NeuroImage 184:293–316.

Kempf F, Kühn AA, Kupsch A, Brücke C, Weise L, Schneider GH, Brown P (2007) Premovement activities in the subthalamic area of patients with Parkinson’s disease and their dependence on task. European Journal of Neuroscience 25:3137–3145.

Knowles T, Adams S, Abeyesekera A, Mancinelli C, Gilmore G, Jog M (2018) Deep brain stimulation of the subthalamic nucleus parameter optimization for vowel acoustics and speech intelligibility in Parkinson’s disease. Journal of Speech, Language, and Hearing Research 61:510–524.

Kuznetsova A, Brockhoff PB, Christensen RHB (2017). lmerTest package: Tests in linear mixed effects models. Journal of Statistical Software 82:1–26.

Lee PS, Weiner GM, Corson D, Kappel J, Chang YF, Suski VR, … & Richardson RM (2018) Outcomes of interventional-MRI versus microelectrode recording-guided subthalamic Deep Brain Stimulation. Frontiers in Neurology 9:241.

Lipski WJ, Alhourani A, Pirnia T, Jones PW, Dastolfo-Hromack C, Helou LB, … Richardson RM (2018) Subthalamic nucleus neurons differentially encode early and late aspects of speech production. Journal of Neuroscience, 38:5620–5631.

Lipski WJ, Wozny TA, Alhourani A, Kondylis ED, Turner RS, Crammond DJ, Richardson RM (2017) Dynamics of human subthalamic neuron phase-locking to motor and sensory cortical oscillations during movement. Journal of Neurophysiology 118:1472–1487.

Lofredi R, Neumann WJ, Bock A, Horn A, Huebl J, Siegert S, … Kühn AA (2018) Dopamine-dependent scaling of subthalamic gamma bursts with movement velocity in patients with Parkinson’s disease. Elife 7:e31895.

Logemann JA, Fisher HB, Boshes B, Blonsky ER (1978) Frequency and co-occurrence of vocal tract dysfunctions in the speech of a large sample of Parkinson patients. J Speech Hear Disord 43:47–57.

Lotte F, Brumberg JS, Brunner P, Gunduz A, Ritaccio AL, Guan C, Schalk G (2015) Electrocorticographic representations of segmental features in continuous speech. Frontiers in Human Neuroscience 9:97.

Lotze M, Seggewies G, Erb M, Grodd W, Birbaumer N (2000) The representation of articulation in the primary sensorimotor cortex. Neuroreport 11: 2985–2989

Meier JD, Aflalo TN, Kastner S, Graziano MS (2008) Complex organization of human primary motor cortex: a high-resolution fMRI study. Journal of Neurophysiology 100:1800–1812.

Monakow HWK, Akert K, Kiinzle H (1978) Projections of the precentral motor cortex and other cortical areas of the frontal lobe to the subthalamic nucleus in the monkey. Exp. Brain Res 33:395–403.

Moore MW, Fiez JA, Tompkins CA (2017) Consonant age-of-acquisition effects in nonword repetition are not articulatory in nature. J Speech Lang Hear Res 60:3198–3212.

Morrison CE, Borod JC, Perrine K, Beric A, Brin MF, Rezai A, Kelly P, Sterio D, Germano I, Weisz D, Olanow CW (2004) Neuropsychological functioning following bilateral subthalamic nucleus stimulation in Parkinson’s disease. Arch Clin Neuropsychol 19:165–181.

Mugler EM, Patton JL, Flint RD, Wright ZA, Schuele SU, Rosenow J, … Slutzky MW (2014) Direct classification of all American English phonemes using signals from functional speech motor cortex. Journal of Neural Engineering 11:035015.

Nadeau SE, Crosson B (1997) Subcortical aphasia. Brain Lang 58:355–402

Nambu A (2011) Somatotopic organization of the primate basal ganglia. Frontiers in Neuroanatomy 5:26.

Nambu A, Takada M, Inase M, Tokuno H (1996) Dual somatotopical representations in the primate subthalamic nucleus: evidence for ordered but reversed body-map transformations from the primary motor cortex and the supplementary motor area. Journal of Neuroscience 16:2671–2683.

Oostenveld R, Fries P, Maris E, Schoffelen J-M (2011) FieldTrip: Open source software for advanced analysis of MEG, EEG, and invasive electrophysiological data. Comput Intell Neurosci 2011:156869.

Penfield W (1954) Mechanisms of voluntary movement. Brain 77:1–17

Penfield W, Boldrey E (1937) Somatic motor and sensory representation in the cerebral cortex of man as studied by electrical stimulation. Brain 60:389–443.

Pulvermüller F, Huss M, Kherif F, del Prado Martin FM, Hauk O, Shtyrov Y (2006) Motor cortex maps articulatory features of speech sounds. Proceedings of the National Academy of Sciences 103:7865–7870.

R Core Team (2018) R: A language and environment for statistical computing. R foundation for statistical computing, Vienna, Austria. URL https://www.R-project.org/

Ramsey NF, Salari E, Aarnoutse EJ, Vansteensel MJ, Bleichner MB, Freudenburg ZV (2017) Decoding spoken phonemes from sensorimotor cortex with high-density ECoG grids. NeuroImage 180: 301–311

Randazzo MJ, Kondylis ED, Alhourani A, Wozny TA, Lipski WJ, Crammond DJ, Richardson RM (2016) Three-dimensional localization of cortical electrodes in deep brain stimulation surgery from intraoperative fluoroscopy. Neuroimage 125:515–521.

Redgrave P, Prescott TJ, Gurney K (1999) The basal ganglia: a vertebrate solution to the selection problem? Neuroscience 89:1009–1023.

Rodriguez-Oroz MC, Rodriguez M, Guridi J, Mewes K, Chockkman V, Vitek J, DeLong MR, Obeso JA (2001) The subthalamic nucleus in Parkinson’s disease: Somatotopic organization and physiological characteristics. Brain 124:1777–1790.

Son RJV, Santen JPV (1997) Strong interaction between factors influencing consonant duration. Fifth European Conference on Speech Communication and Technology.

Starr PA, Theodosopoulos PV, Turner R (2003) Surgery of the subthalamic nucleus: use of movement-related neuronal activity for surgical navigation. Neurosurgery 53:1146–1149.

Tadel F, Baillet S, Mosher JC, Pantazis D, Leahy RM (2011) Brainstorm: A User-Friendly Application for MEG/EEG Analysis. Computational Intelligence and Neuroscience 8.

Takai O, Brown S, Liotti M (2010) Representation of the speech effectors in the human motor cortex: somatotopy or overlap? Brain and language 113:39–44.

Temel Y, Blokland A, Steinbusch HW, Visser-Vandewalle V (2005) The functional role of the subthalamic nucleus in cognitive and limbic circuits. Prog Neurobiol 76:393–413.

Theodosopoulos PV, Marks WJ, Christine C, Starr PA (2003) The locations of movement-related cells in the human subthalamic nucleus in Parkinson’s disease. Movement Disorders 18:791–798.

Umeda N (1977) Consonant duration in American English. The Journal of the Acoustical Society of America 61:846–858.

Wallesch CW, Kornhuber HH, Brunner RJ, Kunz T, Hollerbach B, Suger G (1983) Lesions of the basal ganglia, thalamus, and deep white matter: Differential effects on language functions. Brain and Language 20:286–304.

Walsh B, Smith A (2012) Basic parameters of articulatory movements and acoustics in individuals with Parkinson’s disease. Movement Disorders 27:843–850.

Watson P, Montgomery Jr EB (2006) The relationship of neuronal activity within the sensori-motor region of the subthalamic nucleus to speech. Brain and Language 97:233–240.

Wichmann T, Bergman H, DeLong MR (1994) The primate subthalamic nucleus. I. Functional properties in intact animals. Journal of Neurophysiology 72:494–506.

Witt K, Daniels C, Reiff J, Krack P, Volkmann J, Pinsker MO, Krause M, Tronnier V, Kloss M, Schnitzler A, Wojtecki L, Bötzel K, Danek A, Hilker R, Sturm V, Kupsch A, Karner E, Deuschl G (2008) Neuropsychological and psychiatric changes after deep brain stimulation for Parkinson’s disease: a randomised, multicentre study. Lancet Neurol 7:605–614.

Woolsey CN, Erickson TC, Gilson WE (1979) Localization in somatic sensory and motor areas of human cerebral cortex as determined by direct recording of evoked potentials and electrical stimulation. Journal of Neurosurgery 51:476–506.

